# Conjugal DNA transfer in the maternally inherited symbiont of tsetse flies *Sodalis glossinidius*

**DOI:** 10.1101/2020.06.17.158519

**Authors:** Christopher G. Kendra, Chelsea M. Keller, Roberto E. Bruna, Mauricio H. Pontes

**Affiliations:** Department of Pathology and Laboratory of Medicine, Department of Microbiology and Immunology, Pennsylvania State University College of Medicine, Hershey, PA 17033, USA

**Keywords:** *Sodalis glossinidius*, insect endosymbiont, symbiont, transformation, conjugation, genetic modification, plasmid transfer, transposition, allelic replacement, mutation, paratransgenesis, *Trypanosoma brucei*

## Abstract

Stable associations between insects and bacterial species are widespread in nature. This is the case for many economically important insects, such as tsetse flies. Tsetse flies are the vectors of *Trypanosoma brucei*, the etiological agent of African trypanosomiasis—a zoonotic disease that incurs a high socioeconomic cost in endemic regions. Populations of tsetse flies are often infected with the bacterium *Sodalis glossinidius*. Following infection, *S. glossinidius* establishes a chronic, stable association characterized by vertical (maternal) and horizontal (paternal) modes of transmission. Due to the stable nature of this association, *S. glossinidius* has been long sought as a means for the implementation of *anti-Trypanosoma* paratransgenesis in tsetse flies. However, the lack of tools for the genetic modification of *S. glossinidius* has hindered progress in this area. Here we establish that *S. glossinidius* is amenable to DNA uptake by conjugation. We show that conjugation can be used as a DNA delivery method to conduct forward and reverse genetic experiments in this bacterium. This study serves as an important step in the development of genetic tools for *S. glossinidius*. The methods highlighted here should guide the implementation of genetics for the study of the tsetse-*Sodalis* association and the evaluation of *S. glossinidius*-based tsetse fly paratransgenesis strategies.

**Importance:** Tsetse flies are the insect vectors of *T. brucei*, the causative agent of African sleeping sickness—a zoonotic disease that inflicts a substantial economic cost to a broad region of sub-Saharan Africa. Notably, tsetse flies can be infected with the bacterium *S. glossinidius* to establish an asymptomatic chronic infection. This infection can be inherited by future generations of tsetse flies allowing *S. glossinidius* to spread and persist within populations. To this effect, *S. glossinidius* has been considered as a potential expression platform to create flies which reduce *T. brucei* stasis and lower overall parasite transmission to humans and animals. However, the efficient genetic manipulation of *S. glossinidius* has remained a technical challenge due to its complex growth requirements and uncharacterized physiology. Here we exploit a natural mechanism of DNA transfer among bacteria and develop an efficient technique to genetically manipulate *S. glossinidius* for future studies in reducing trypanosome transmission.

## Introduction

African trypanosomiasis or sleeping sickness is a zoonotic disease caused by the parasitic protozoa *Trypanosoma brucei. Trypanosoma brucei* is transmitted by tsetse flies (*Glossina* spp.; Diptera: *Glossinidae*), viviparous insects that feed exclusively on vertebrate blood (1, 2). In addition to *T. brucei*, natural populations of tsetse flies are often infected with strains of the Gram negative bacterium *Sodalis glossinidius* (3–7). The establishment of *S. glossinidius* infection leads to a stable association, where the bacterium colonizes a number of tsetse fly tissues, including the salivary glands inhabited by *T. brucei*, without imposing a measurable burden to the flies (4, 8–12). Importantly, while *S. glossinidius* undergoes a predominantly maternal mode of transmission, being passed from mother to offspring during gestation (3, 8–12), this bacterium is also capable of paternal transmission during copulation (13), a phenomenon that may facilitate its colonization and spread within uninfected tsetse populations. Due to these particular characteristics, *S. glossinidius* has emerged as an attractive candidate for the implementation of tsetse fly paratransgenesis—a bioremediation strategy where bacteria capable of colonizing tsetse populations are used to express traits that inhibit *Trypanosoma* transmission (14–19).

The development of *Sodalis*-based paratransgenesis relies on the ability to genetically modify this bacterium. Although *S. glossinidius* has been isolated in axenic culture (4) and its genome has been sequenced (20), this bacterium has been proven refractory to artificial DNA transformation techniques. To date, two artificial transformation methods have been employed to introduce of exogenous plasmid DNA into *S. glossinidius:* heat-shock transformation and electroporation (8, 10, 13, 14, 16, 17, 21–25). While these transformation procedures were originally developed for *Escherichia coli* and popularized by the use of this organism as the workhorse of molecular biology (26, 27), they have proven to be both unreliable and inefficient as DNA delivery methods for *S. glossinidius*. Additionally, the use of natural DNA transfer methods such as conjugation have been hindered by the complex nutritional requirements and slow growth rate of *S. glossinidius* (28), which undermines strategies to counter-select donor bacteria following DNA transfer.

Here, we identify growth conditions enabling the counter-selection of *E. coli* donor strains and demonstrate that *S. glossinidius* is amenable to uptake of DNA by conjugation (Fig. 1). We show that *S. glossinidius* exogenous DNA recipients (transconjugants) can be readily and reproducibly recovered after biparental mating with these *E. coli* donor strains. We use this technique for the implementation of forward genetic analysis through the generation of random transpositions with a Himar1 mariner and mini-Tn*5* transposition systems, and reverse genetics by insertionally inactivating a number of chromosomal genes using suicide vectors. This work establishes conjugation as a reliable DNA delivery method for the genetic manipulation of *S. glossinidius* and will greatly facilitate the study of this bacterium and the evaluation of methods for tsetse paratransgenesis.

**Fig. 1.**
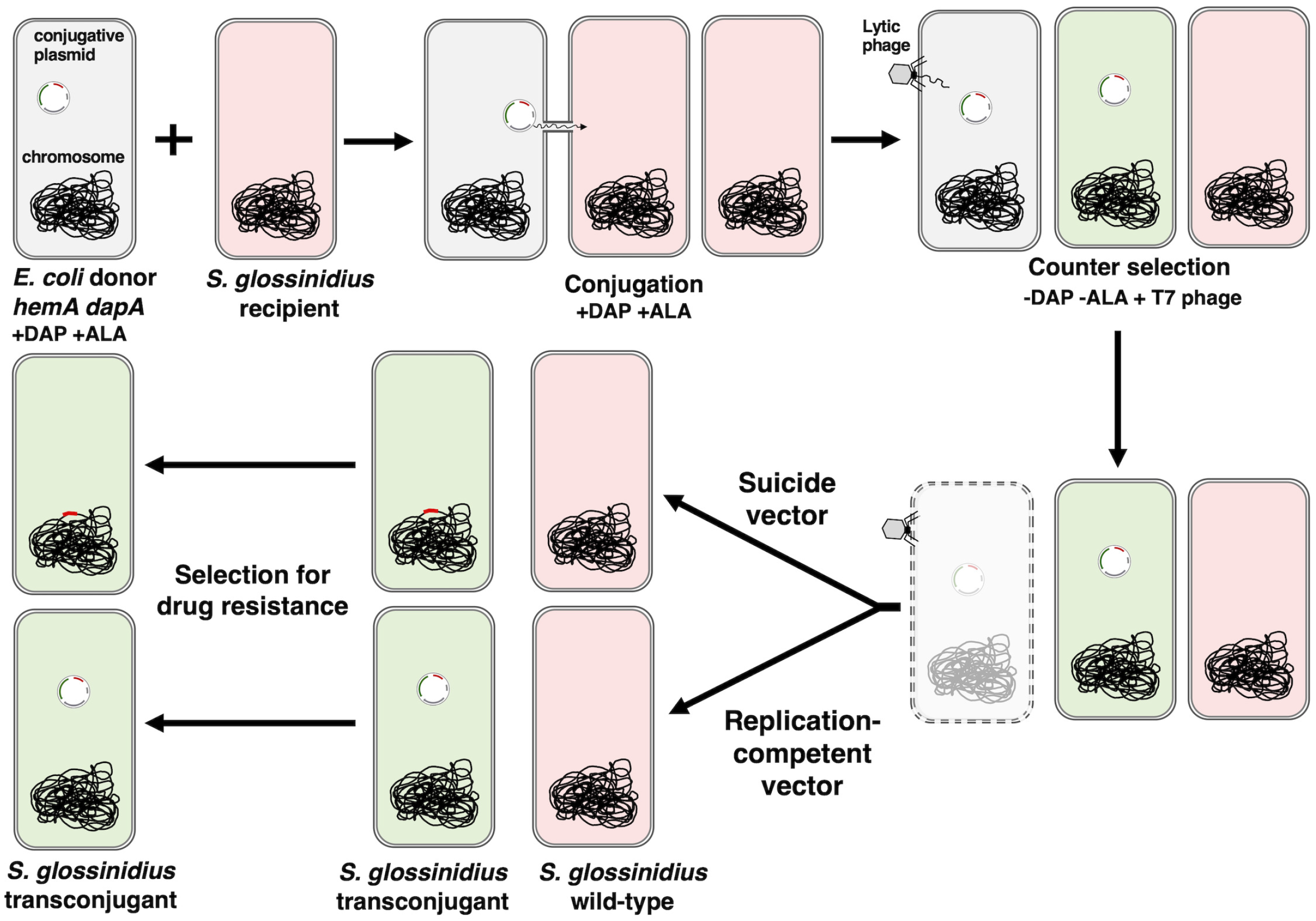
Representative workflow of the conjugation procedure developed for *Sodalis glossinidius*. An *E. coli hemA dapA* donor strain (gray) is mixed and grown with *S. glossinidius* (red) in the presence of DAP and ALA. Following the transfer of a mobilizable genetic element from donor to recipient cells, the majority of *E. coli* cells are eliminated by exposing the mixed culture to the lytic bacteriophage T7. The cell mixture is washed and plated on medium lacking DAP and ALA. The presence of a selective agent on the plate selects for *S. glossinidsius* transconjugants (green cells that received DNA from the *E. coli hemA dapA* donor).

## Results

### Prolonged incubation of *Escherichia coli dapA hemA* donor strain on rich medium lacking diaminopimelic acid and δ-aminolevulinic acid gives rise to suppressor mutants that no longer require these nutrients

During conjugation, donor and recipient cells must come into close physical proximity to enable DNA transfer through a pilus. Subsequently, recipient cells which have received DNA (transconjugants) are recovered on plates containing solid medium that restricts the growth of donor and recipient cells that did not receive the desired DNA molecule. Transconjugants are positively selected based on the presence of a genetic marker within the transferred DNA (i.e. antibiotic resistant gene). However, because donor cells retain a copy of the transferred DNA, they have to be eliminated by other means. Classically, this is achieved through nutritional-based auxotrophy counter selections, where the growth of the donor is hindered by plating the conjugation mixture on defined medium lacking a nutrient synthesized by the recipient, but not the donor (e.g. a particular amino acid). However, amino acid-auxotrophy-based strategies cannot be used to counter-select donor cells in conjugation mixtures with *S. glossinidius*. This is because *S. glossinidius* is a slow-growing microaerophilic bacterium with complex nutritional requirements, and defined solid media recipes that support the formation of colonies are currently unavailable. To overcome this hurdle, we sought to exploit well-characterized *E. coli* donor strains containing alternative auxotrophies that can be used for counter-selection on complex media.

In *E. coli*, the *dapA* gene encodes a 4-hydroxy-tetrahydrodipicolinate synthase and the *hemA* gene encodes a glutamyl-tRNA reductase. These enzymes are required for the biosynthesis of peptidoglycan and heme, respectively. While mutations in *dapA* give rise to a requirement for diaminopimelic acid (DAP), mutations in *hemA* create a requirement for δ-aminolevulinic acid (ALA) or heme, respectively (Fig. 2A) (29, 30). As DAP and ALA are usually not present in complex microbial medium components, *E. coli* donor strains containing these mutations are often used to select transconjugants on rich medium such as Luria Bertani (LB) (31, 32). *Sodalis glossinidius* forms colonies 5 to 10 days following plating on rich media, such as brain heart infusion-blood (BHIB) agar. Therefore, we sought to determine if *E. coli* donor strains containing mutations in *dapA* and/or *hemA* were able to grow on BHIB agar. *Escherichia coli dapA* and *hemA* strains were streaked on BHIB agar alongside with *S. glossinidius* and incubated for 8 days under microaerophilic conditions. Following incubation, *S. glossinidius* formed small colonies as expected (Fig. 2B). By contrast, *dapA* and *hemA* strains displayed residual growth at the inoculation sites on the plates (Fig. 2B). Control BHIB plates supplemented with DAP supported the growth the *E. coli dapA* strain, which formed large colonies following 8 days of incubation (Fig. 2B).

**Fig. 2.**
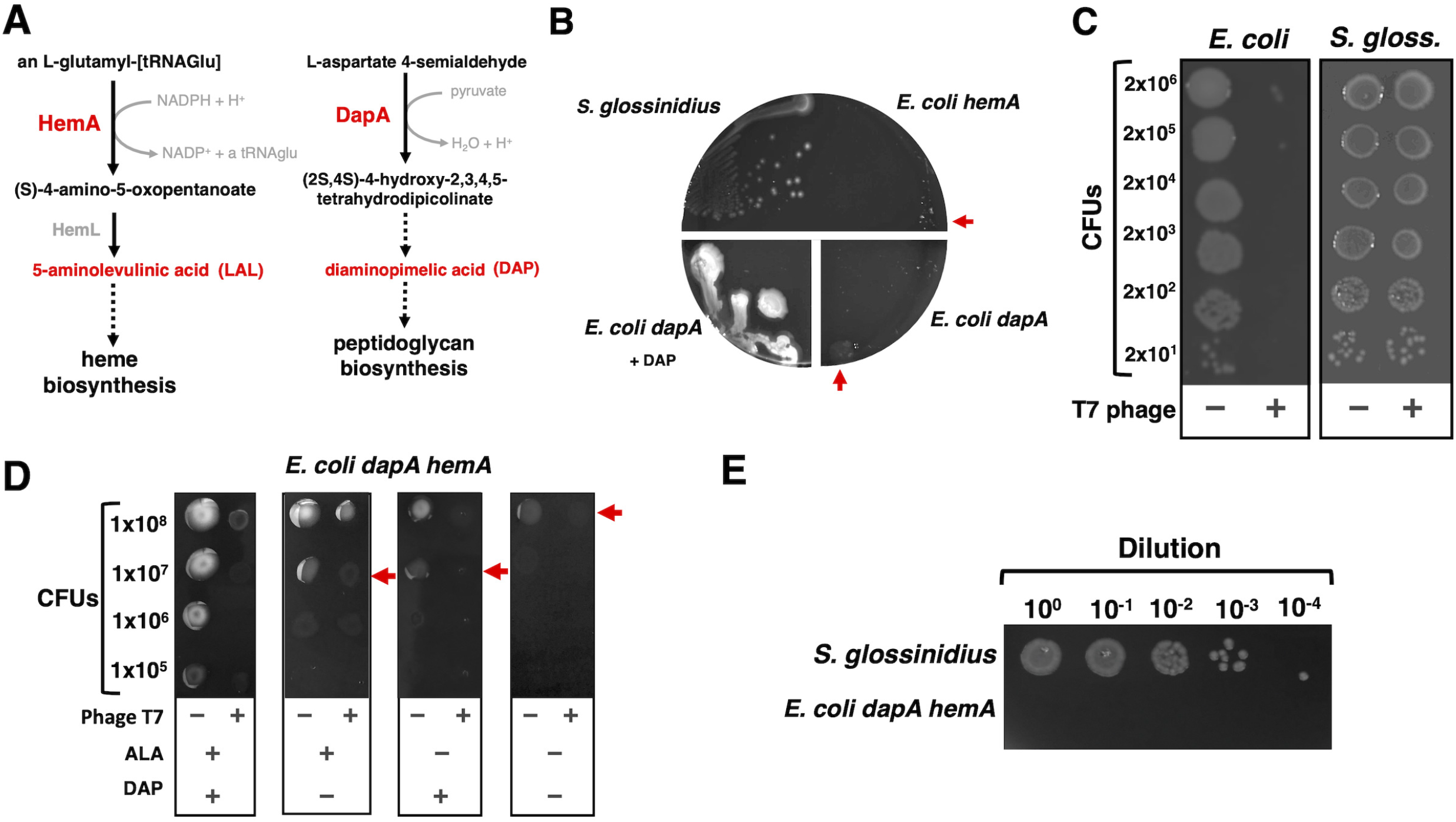
Counterselection of *E. coli hemA dapA* donor on BHIB agar. (A) Schematics depicting the reactions catalyzed by HemA (left-hand side) and DapA (right-hand side), enzymes required for the biosynthesis of heme and peptidoglycan respectively. Complementation of growth medium with the metabolic intermediates depicted in red are used to support the growth of *hemA* and *dapA* mutants. (B) Growth of wild-type *S. glossinidius, E. coli hemA* (MP1182) and *E. coli dapA* (BW29427) on BHIB agar lacking or containing DAP. Plates were photographed after 8 days of incubation at 27°C under microaerophilic conditions. Red arrows indicate residual growth. (C) Growth of wild-type *E. coli* (MG1655) and *S. glossinidius* following 120 min incubation in 10 mM MgCl_2_ or 10 mM MgCl2 containing phage T7. Cell suspensions were diluted and 5 μL were spotted on plates. *Escherichia coli* was incubated at 37°C on LB for 16h. *Sodalis glossinidius* was incubated under microaerophilic conditions at 27°C on BHIB for 8 days. (D) Growth of *E. coli dapA hemA* (MP1554) on BHIB agar with various combinations of ALA and DAP. Cells were incubated for 120 min in 10 mM MgCl2 or 10 mM MgCl2 containing phage T7. Cultures were diluted and 5 μL were plated on BHIB agar. Plates were incubated at 37°C for 16h. Red arrows indicate residual growth. (E) Growth of *S. glossinidius* and *E. coli dapA hemA* (MP1554) on BHIB agar lacking ALA and DAP. Bacteria were grown separately on plates in a mock conjugation experiment, subsequently exposed to phage T7, washed, diluted and spotted on BHIB as described in (C). Plates were incubated for 8 days at 27°C under microaerophilic conditions. Images depict representative plates of at least 3 independent experiments.

The aforementioned results suggested that it might be possible to counter select a *dapA* or *hemA E. coli* donor strain on BHIB agar following conjugation with *S. glossinidius*. We therefore attempted to recover *S. glossinidius* transconjugants under a number of mating conditions. We found that *E. coli dapA* suppressor mutants that are able to grow in the absence of DAP emerge at high frequency following 5 or 16 h of mating, where strains are mixed at ratios of 50 *Sodalis* to 1 *E.coli* or 2,500 *Sodalis* to 1 *E.coli*, respectively (Fig. S1A). Indeed, even in an *E. coli* donor strain containing both *dapA* and *hemA* mutations, suppressor mutants that do not require DAP and ALA emerged at a relatively high frequency following 8 days of incubation on BHIB agar (approximately 2×10^−7^ CFUs; Fig. S1B). Together, these results indicate that the introduction of *dapA* and *hemA* mutations in an *E. coli* donor strain can be used as part of a counter-selection strategy, but are not sufficient to retrieve *S. glossinidius* transconjugants.

### *dapA hemA* donor suppressors can be eliminated using an *E. coli*-specific lytic bacteriophage

T7 is a lytic bacteriophage (phage) with a narrow host range. This phage typically infects certain *E. coli* and closely related *Shigella* strains, as well as certain *Yersinia* strains (33). Given the specificity of T7, we wondered if we could use this phage to target *E. coli* cells in conjugation mixtures with *S. glossinidius*. Therefore, we sought to determine if *S. glossinidius* was immune to killing by phage T7. We established that despite causing a decrease of five orders of magnitude in the number of colony-forming units (CFUs) in cultures of the *E. coli* donor strain, exposure to phage T7 did not decrease CFU counts in *S. glossinidius* cultures, indicating that the later bacterium is immune to T7 killing (Fig. 2C).

Given these results, we decided to examine the effect of phage T7 on the emergence of *E. coli dapA hemA* suppressors that can grow in the absence of DAP, ALA, or both. We plated the *E. coli dapA hemA* donor on BHIB agar in the presence or absence of DAP, ALA, and/or phage T7. Consistent with previous results (Fig. 2B and S1B), removal of either or both DAP and ALA caused a decrease in cell survival of the *E. coli* donor (Fig. 2D). The presence of phage T7 alone also decreased the survival of the *E. coli* donor under all conditions tested (Fig. 2D). In the absence of DAP and ALA, phage T7 lowered the number of donor cells by over nine orders of magnitude, effectively preventing the emergence of *E. coli dapA hemA* suppressors that can grow in the absence of DAP and ALA (Fig. 2D and S1B). Importantly, after population expansion of the *E. coli* donor for 16 h in a mock conjugation experiment, exposure to phage T7 was sufficient to prevent the emergence of *E. coli dapA hemA* suppressors within the time window permitting *S. glossinidius* to form colonies (i.e. 8 days; Fig. 2E). Together, these results suggested that *S. glossinidius* cells can be isolated from conjugation mixtures with an *E. coli dapA hemA* donor following exposure to T7 phage.

### Conjugation of transposition systems for random mutagenesis of *Sodalis glossinidius*

Transposable elements have played a pivotal role in the development of forward genetics studies in bacterial species (34, 35), and have been previously used in studies of *S. glossinidius* (22). We therefore attempted to use conjugation for the delivery of stable transposition systems encoded within mobilizable suicide vectors into this bacterium. Following conjugation, *S. glossinidius* transconjugants were readily recovered by selecting for the antibiotic markers encoded within each transposon (Fig. 3A and B).

**Fig. 3.**
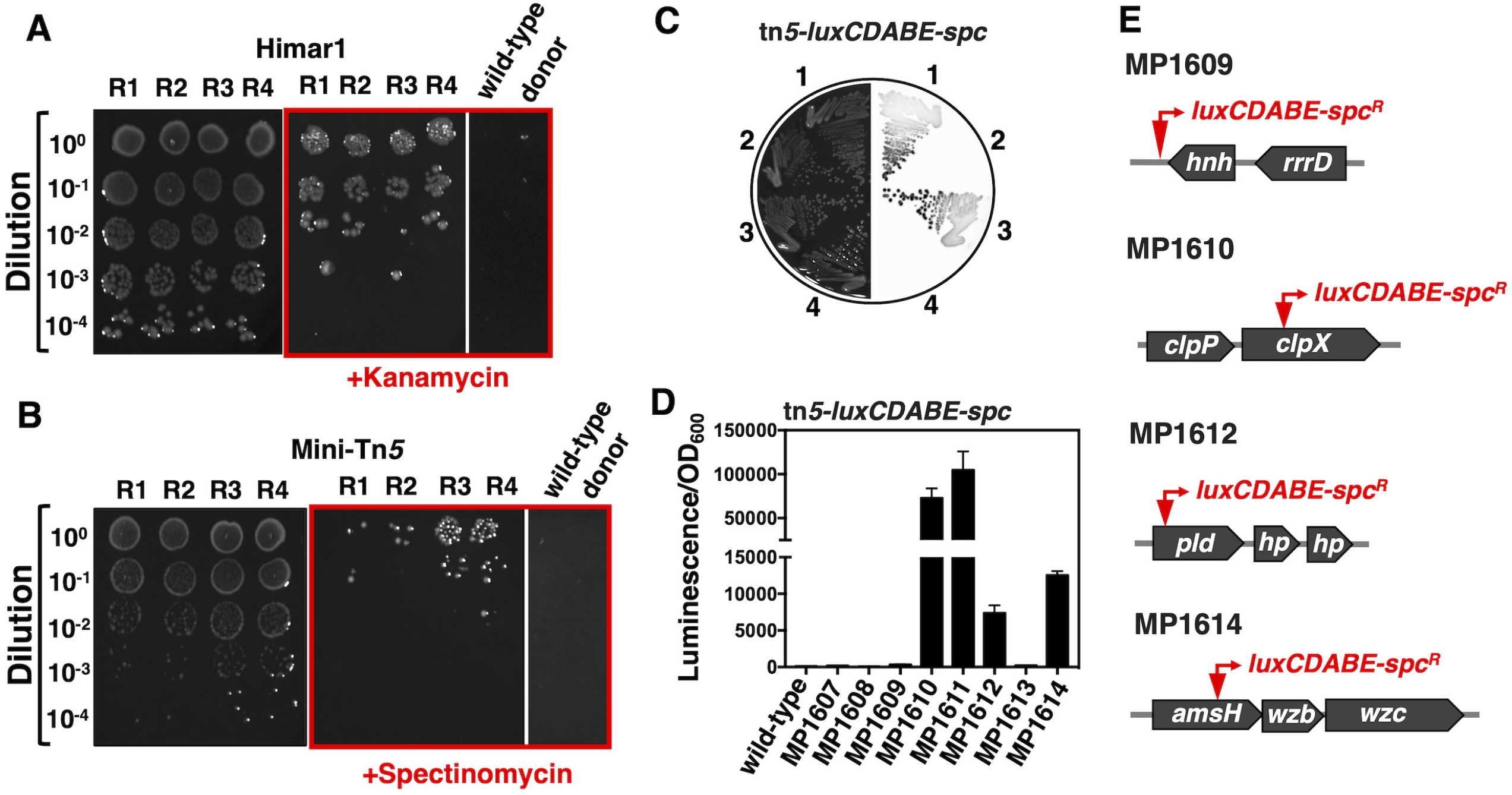
Transposition mutagenesis in *S. glossinidius*. (A) Serial dilutions of conjugation mixtures of *S. glossinidius* and *E. coli dapA hemA* (MP1554) harboring the suicide vector encoding a Mariner transposon system (Himar1), pMarC9-R6k. 5 μL of cell suspension were spotted on BHIB agar (left-hand-side panel) or BHIB agar supplemented with kanamycin (middle panel). Individually grown conjugation partners, *S. glossinidius* and *E. coli dapA hemA* (MP1554) pMarC9-R6k, were also spotted on BHIB agar supplemented with kanamycin (right-hand-side panel). The red square indicates plates containing kanamycin. (B) Serial dilutions of conjugation mixtures of *S. glossinidius* and *E. coli dapA hemA* (MP1554) harboring the suicide vector encoding the Tn*5*-based promoter-probe transposition system, pUTmini-Tn*5*-*luxCDABE*-Spc. 5 μL of cell suspension were spotted on BHIB agar (left-hand-side panel) or BHIB agar supplemented with spectinomycin (middle panel). Individually grown *S. glossinidius* and *E. coli dapA hemA* (MP1554) pUTmini-Tn*5*-*luxCDABE*-Spc were spotted on BHIB agar supplemented with spectinomycin (right-hand-side panel). Red square indicates plates containing spectinomycin. (C) Transconjugants obtained in a conjugation experiment described in (B) were purified on BHIB agar supplemented with spectinomycin. Luminescence signals of four distinct clones are depicted on the righth-and-side of the figure. Plates were incubated for 8 days at 27°C under microaerophilic conditions. Images depict representative plates of at least 3 independent experiments. (D) Quantification of luminescence signals derived from selected *S. glossinidius* mini-*Tn5-luxCDABE-spc^R^* transconjugants obtained as described in (B). Error bar represent standard deviations from three technical replicates. (E) Schematic illustration depicting locations of *mini-Tn5-luxCDABE-spc^R^* transposition insertions in selected *S. glossinidius* clones—*hnh* (SGGMMB4_03814), *clpX* (SGGMMB4_01523), *pld* (SGGMMB4_05728) and *amsH* (SGGMMB4_02193).

A number of controls indicated that these *S. glossinidius* cells were true transconjugants resulting from random transposition events, originating from the mobilized suicide vector. First, no antibiotic resistant clones were recovered from *S. glossinidius* cells which were not conjugated with the *E. coli* donor. Hence, the emergence of antibiotic resistance was linked to a physical interaction with the donor strain (Fig. 3A and B). Second, antibiotic resistant clones of *S. glossinidius* remained sensitive to ampicillin, indicating that they did not retain the suicide vector, either as an autonomous replicating episome or as a vector integrated into the chromosome (Table 1). Third, conjugation experiments involving promoter-probe transposition systems, such as the *Tn5-luxCDABE-Spc* (36), yielded a population of antibiotic resistant *S. glossinidius* clones displaying heterogeneous reporter-gene expression (Fig. 3C and D). Thus, the recovered transconjugant clones emerged from distinct transposition events (Fig. 3C and D). Consistent with these observations, mapping putative transposition events in some of these clones revealed transposon insertions into distinct chromosomal locations (Fig. 3E). Importantly, while the conjugation efficiency varied with particular transposition systems, transconjugants were reproducibly recovered at high frequency (1.40 x 10^−3^ to 1.84 x 10^−2^) (Table 1 and Table S1). Together, these results demonstrate that conjugation can be reliably used to deliver transposition systems into *S. glossinidius*.

**Table 1.** Summary of matting experiments

| Plasmid | Origin of replication | Positive selection |  | Conjugation efficiency | Plasmid retention (Amp<sup>R</sup>) |
| --- | --- | --- | --- | --- | --- |
|  |  | Transposition or insertion | Plasmid |  |  |
| pFAJ1815 | R6K $\gamma$ | Kanamycin | Ampicillin | 1.80E-03 (N=14) | 0/24 |
| pUT-miniTn5-lux-Km2 | R6K $\gamma$ | Kanamycin | Ampicillin | 6.82E-03 (N= 21) | 0/24 |
| pUT-Tn5-GFP | R6K $\gamma$ | Kanamycin | Ampicillin | 5.56E-03 (N=12) | N/D |
| pMarC9-R6K | R6K $\gamma$ | Kanamycin | Ampicillin | 1.84E-02 (N=5) | N/D |
| pUT-miniTn5-lux-Sp | R6K $\gamma$ | Spectinomycin | Ampicillin | 1.40E-03 (N=8) | 0/21 |

Amp<sup>R</sup> – ampicillin resistance. N/D – not determined.

### Conjugation of suicide vectors for targeted gene disruption in *Sodalis glossinidius*

We tested whether we could use conjugation for the delivery of replication-deficient suicide plasmids designed for targeted gene disruption. In contrast to transposition, this reverse genetic strategy relies on homologous recombination functions encoded by the host bacterium (37). In its simplest form, insertional disruptions can be generated through single homologous recombination events between the target gene and a homologous fragment cloned in a suicide vector—i.e. a Campbell-like integration (Fig. 4A). We employed this strategy to target the transcriptional regulators encoded by *S. glossinidius cpxR* and *ompR* genes. Following conjugation, we were able to recover antibiotic resistant *S. glossinidius* clones which, upon polymerase chain reaction (PCR) analyses, were shown to harbor plasmid insertions in the expected chromosomal locations (Fig. 4B and C). Taken together, these results demonstrate that conjugation can be used for the delivery of suicide vectors for targeted gene disruption in *S. glossinidius*.

**Fig. 4.**
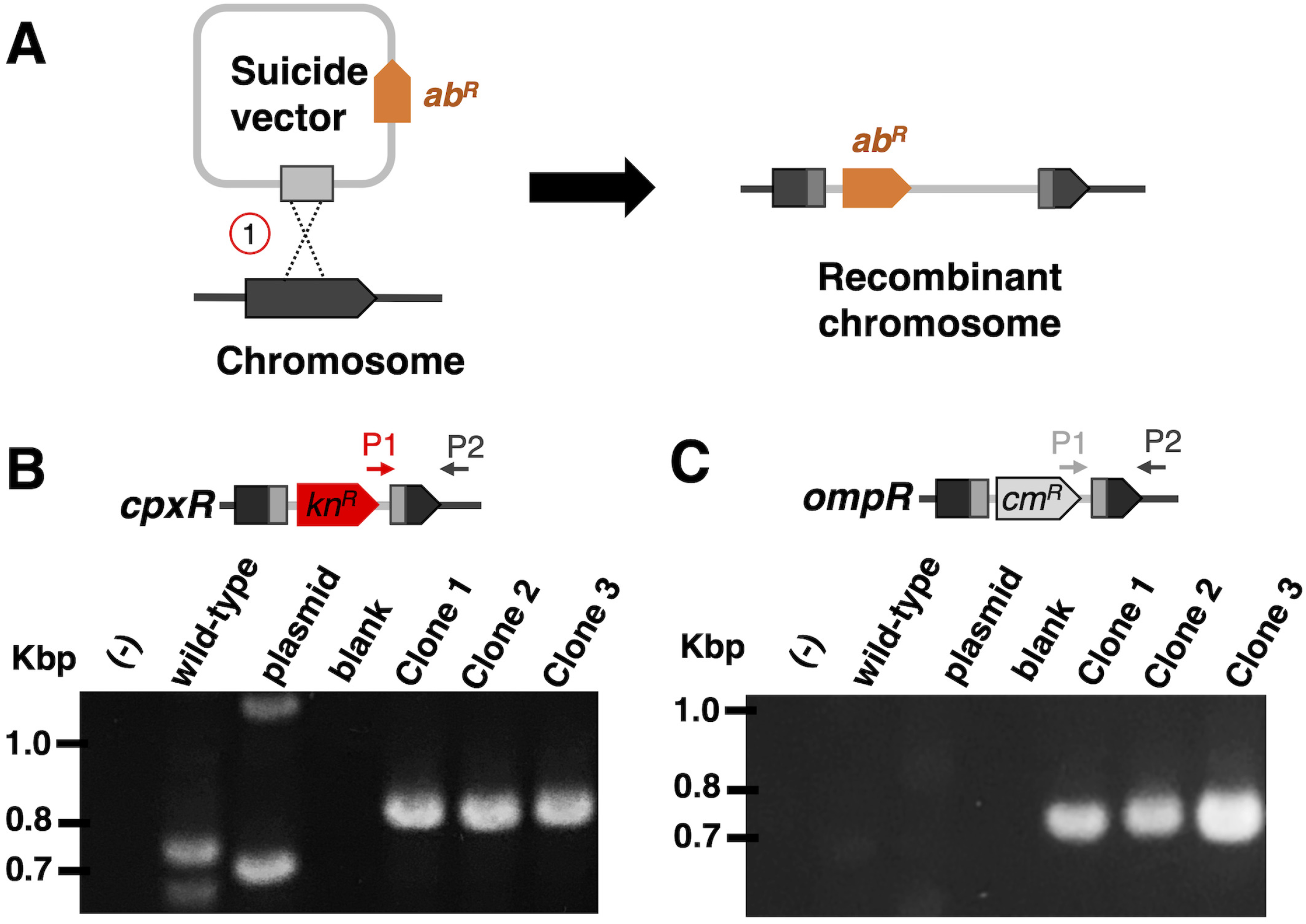
Gene targeting in *S. glossinidius* by insertional inactivation. (A) Schematic depicting the integration of a suicide vector harboring an antibiotic resistant marker (*ab^R^*) into a specific chromosomal gene by a homologous recombination event (1). (B and C) PCR confirmation of integration events disrupting *S. glossinidius cpxR* and *ompR* homologs.

## Discussion

In the current study, we established conditions permitting the counter-selection of *E. coli* on BHIB agar. We use these conditions to hinder the growth of *E. coli* DNA donor strains following mating, and demonstrate that the slow-growing, fastidious bacterium *Sodalis glossinidius* is receptive to DNA transfer by conjugation. We employed conjugation to perform random transposition and targeted mutagenesis in *S. glossinidius*, effectively implementing efficient methods to carry out forward and reverse genetics. Similar procedures can be developed for the delivery of any mobilizable genetic elements harboring an origin of transfer (*oriT*) to this bacterium. These include replication-competent plasmids or other episomes encoding an array of functions, such as targeted transposition (38) and advanced genome editing CRISPR-based systems (39). The basic genetic modifications performed in this study serve as a proof of concept, and should greatly facilitate the development and implementation of *S. glossinidius*-based paratransgenesis in tsetse flies. From a broad perspective, this study highlights the application of conjugation as a DNA delivery mechanism for *S. glossinidius* and potentially other fastidious bacterial species with undeveloped genetic methods. These include many insect-associated bacteria such as *Candidatus* Arsenophonus arthropodicus, *Candidatus* Arsenophonus triatominarum, *Spiroplasma poulsonii, Hamiltonella defensa* and *Serratia symbiotica* (40–44).

## Material and Methods

### Microbial strains, phages, plasmids and growth conditions

Microbial strains, phages and plasmids used in this study are presented in Table S2. Unless indicated, all *E. coli* strains were propagated at 37°C or 30°C in Luria Bertani (LB) broth or agar (1.5% w/v). *Sodalis glossinidius* was grown at 27°C in brain-heart infusion broth supplemented with 10 mM MgCl_2_ (BHI) or on brain-heart infusion agar (1.2% w/v) supplemented with 10% defibrinated horse blood and with 10 mM MgCl_2_ (BHIB). Growth of *Sodalis glossinidius* on BHIB was carried out under microaerophilic conditions, which was achieved either using BD GasPak EZ Campy Gas Generating sachets or a gas mixture (5% oxygen 95% CO2). For both *E. coli* and *S. glossinidius*, growth in liquid medium was carried out with aeration (250 rpm). When required, medium was supplemented with ampicillin (100 μg/mL), chloramphenicol (20 μg/mL for *E. coli* and 10 μg/mL for *S. glossinidius*), kanamycin (50 μg/mL for *E. coli* and 25 μg/mL for *S. glossinidius*), spectinomycin (100 μg/mL for *E. coli* and 30 μg/mL for *S. glossinidius*), gentamycin (18 μg/mL for *E. coli* and 9 μg/mL for *S. glossinidius*), δ-aminolevulinic acid (ALA, 100 μg/mL), diaminopimelic acid (DAP, 60 μg/mL). Anhydrotetracycline was used at 0.2 μg/mL.

### Construction of suicide vectors for targeted gene disruption

Oligonucleotide sequences used in this study are presented in Table S3. Phusion® High-Fidelity DNA Polymerase (New England BioLabs) was used in PCR with *S. glossinidius* genomic DNA or plasmids pKD3 or pKD4 (45). While *S. glossinidius* genomic DNA was used for the amplification of fragments flanking genes targeted for gene disruption (*cpxR* and *ompA*), plasmids were used for the amplification of chloramphenicol, kanamycin or gentamycin resistant genes. Replacement alleles were assembled in the backbone of suicide vector pAOJ15 (46), previously digested with BamHI and EcoRI, using NEBuilder® HiFi DNA Assembly (New England BioLabs). Assembly reactions were transformed into *E. coli* cells, and transformants were selected on LB plates containing either ampicillin and chloramphenicol or kanamycin. The integrity of constructs was verified by PCR using primers flanking the ligation points between different fragments within each plasmid.

### Construction of *hemA Escherichia coli* strains

Oligonucleotide sequences used in this study are presented in Table S3. *Escherichia coli* BW29427 (also known as WM3064) or S17-1 harboring plasmid pSIM6 (47) were grown overnight in LB medium supplemented with 100 μg/mL of ampicillin and 60 μg/mL of DAP at 30°C and 250 rpm. Cells were diluted (1:100) in 30 mL of the same medium and grown for approximately 2.5 h (OD_600_ ~0.35-0.4). The cultures were then grown in a water bath for 30 min at 42°C and 250 rpm (final OD_600_ ~0.6-0.8). Cells were immediately transferred to a 50 mL conical tube, collected by centrifugation (4°C at 7,000 rpm for 2.5 min) and resuspended in 40 mL of ice-cold dH_2_O. Cells were collected again by centrifugation and this washing procedure was repeated a second time. Finally, cells were resuspended in 150 μL of ice-cold dH_2_O. Homologous recombination was obtained by electroporating 70 μL of cell suspension with 10 μL of the purified PCR product, generated with primers 212 and 213 and plasmid pKD3 as the template (45). Recombinants were recovered on LB plates supplemented with 20 μg/mL of chloramphenicol, 60 μg/mL of DAP and 100 μg/mL ALA. Chloramphenicol clones were screened for the inability to grow on LB in the absence of ALA.

### Transposition mapping

Mapping of *mini-Tn5-luxCDABE-spc^R^* (36) transposition insertions were carried out as described (48).

### Preparation of bacteriophage T7 solutions

*Escherichia coli* MG1655 cultures were grown overnight in LB broth at 37°C and 250 rpms. One mL of overnight cultures were used to inoculate 100 mL of fresh LB broth. Cultures were allowed to grow for 2 h at 37°C and 250 rpms to an OD_600_ ~0.3-0.4, and then infected with bacteriophage T7 lysate. After 3 to 4 h of lysis, cells were transferred to conical tubes and 1/1000 volume of chloroform was added to each tube. Tubes were vortexed for 1 minute, cell debris were pelleted by centrifugation (7,000 rpm for 2.5 min at room temperature), and the lysate was filtered through a 0.22 μm polyethersulfone membrane filter. Bacteriophage lysate was concentrated using an Amico® Ultra-15 centrifugal filter (Millipore) and LB broth was replaced with a solution of 10 mM MgCl2. Lysates were sterilized by filtration through a 0.22 μm polyethersulfone membrane and stored at 4°C.

### Bacteriophage T7 killing assay

Cultures of *E. coli* MG1655 and *S. glossinidius* were exposed to T7 phage lysate for 30 minutes. Cells were collected by centrifugation, resuspended in fresh medium, diluted and spotted on agar plates. For *E. coli*, plates were incubated for 16 h at 37°C; for *S. glossinidius*, plates were incubated for 8-10 days at 27°C under microaerophilic conditions.

### Conjugation conditions

*Escherichia coli* donor strains were grown overnight (18h) in LB broth supplemented with 100 μg/mL of ampicillin, 60 μg/mL of DAP and/or 100 μg/mL ALA, at 37°C and 250 rpms. Cultures were diluted 1:100 into 3 mL of fresh medium lacking ampicillin and grown for 2 h, to an optical density (OD_600_) ~0.15-0.3). *Sodalis glossinidius* was grown in 150-250 mL of BHI broth to an OD_600_ ~0.4-1.0. Cells were then collected by centrifugation (5 minutes at 7,000 rpms at room temperature) and resuspended in BHI broth to a final an OD_600_ ~10 to 20. *Escherichia coli* donor and *S. glossinidius* recipient were mixed (20 μL donor solution: 200 μL recipient solution) and 50 μL aliquots were spotted on a 0.45 μm filter paper resting on BHIB agar supplemented with DAP and/or ALA. Control mating, containing only donor or recipient cells, were set up alongside, and plates were incubated for 12-16h at 27°C under microaerophilic conditions. Cells on filters were resuspended in a solution of bacteriophage T7 and incubated for 2 h at room temperature. The cell mixture was collected by centrifugation (5 minutes at 7,000 rpms at room temperature) and washed 3 times with phosphate-buffered saline (PBS) to remove residual DAP and ALA. Cell pellets were then resuspended in T7 solution, diluted and plated on BHIB agar with the appropriate antibiotics and 0.2 μg/mL of anhydrotetracycline.

### Quantification of bioluminescence in bacterial cultures

Light production by individual clones grown in BHI broth were measured using a SpectraMax i3x plate reader (Molecular Probes). Luminescence signals were normalized by optical densities of the cultures (absorbance at 600 nm).

### Image Acquisition, Analysis and Manipulation

DNA agarose gel eletrophoresis and light production by *S. glossinidsius* colonies were detected using an Amersham Imager 680 (GE Healthcare). When oversaturated, the intensity of signals in images were adjusted across the entire images using Preview (Apple).

## Acknowledgement

We would like to thank Serap Aksoy (Yale University) for kindly providing us with a culture of *Sodalis glossinidius*, and Kenneth Keiler (Penn State University) for providing us plasmid pUT-Tn5-GFP. MHP is supported by grant AI148774 from the National Institutes of Health and startup funds from The Pennsylvania State University College of Medicine.

## Supplemental Material

Figure S1.

Table S1.

Table S2.

Table S3.

---

**Fig. S1.**
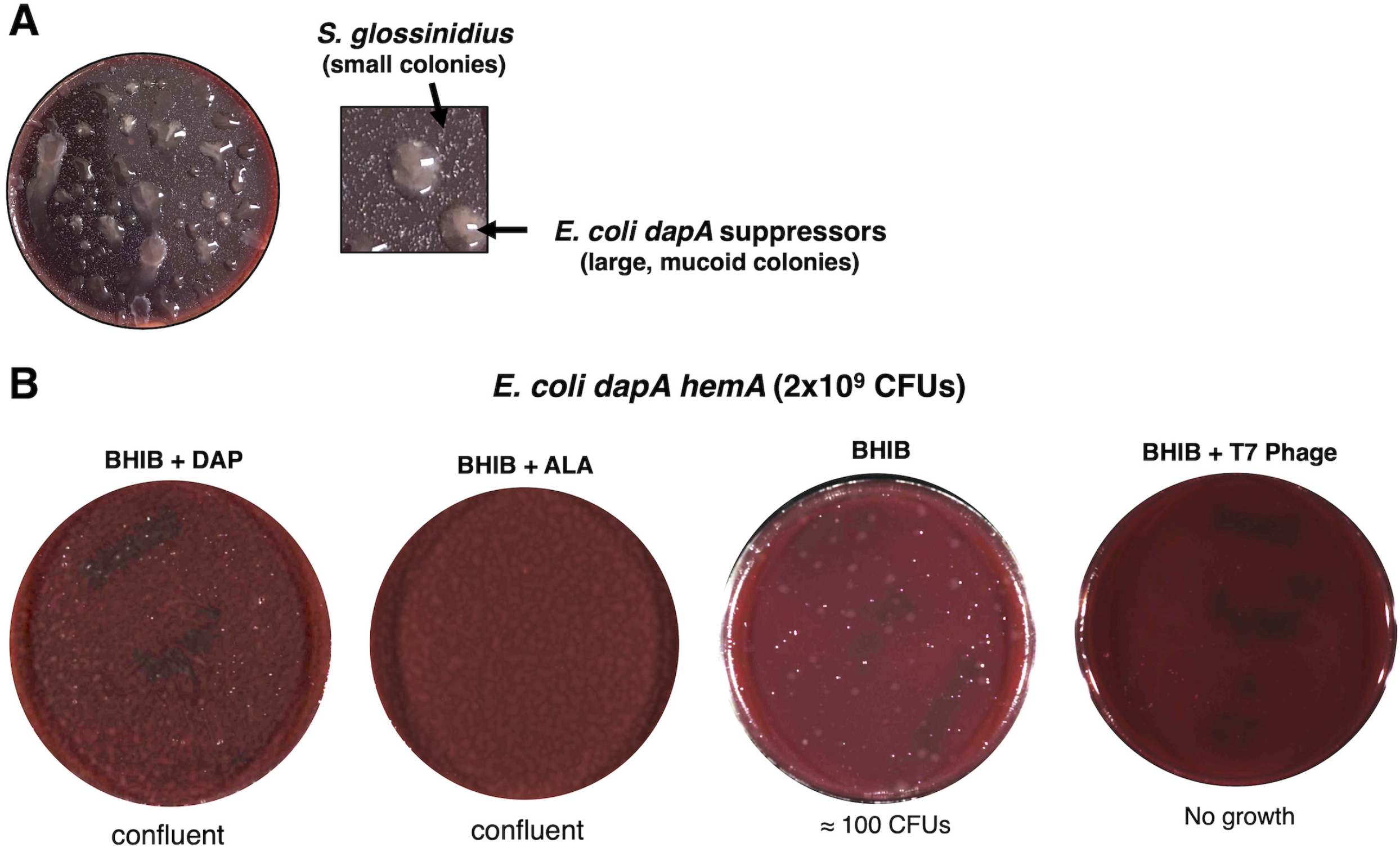
Growth of *E. coli dapA* and *dapA hemA* strains on BHIB agar. (A) A representative plate of an *E. coli dapA* and *S. glossinidius* conjugation mixture grown for 8 days, at 27°C on BHIB agar under microaerophilic conditions. The square highlights a magnified portion of the plate depicting small *S. glossinidius* colonies and large mucoid *E. coli dapA* suppressor colonies that are able to grow in the absence of DAP (29). (B) Representative BHIB agar plates seeded with 2 x 10^9^ CFUs of *E. coli dapA hemA*. Where indicated, plates were supplemented with DAP or ALA, or cells were pre-treated with bacteriophage T7 lysate. Plates were incubated for 7 days, at 27°C and under microaerophilic conditions. Plates displaying confluent growth were imaged following 2 days of incubation, when growth became apparent.

**Table S1.**
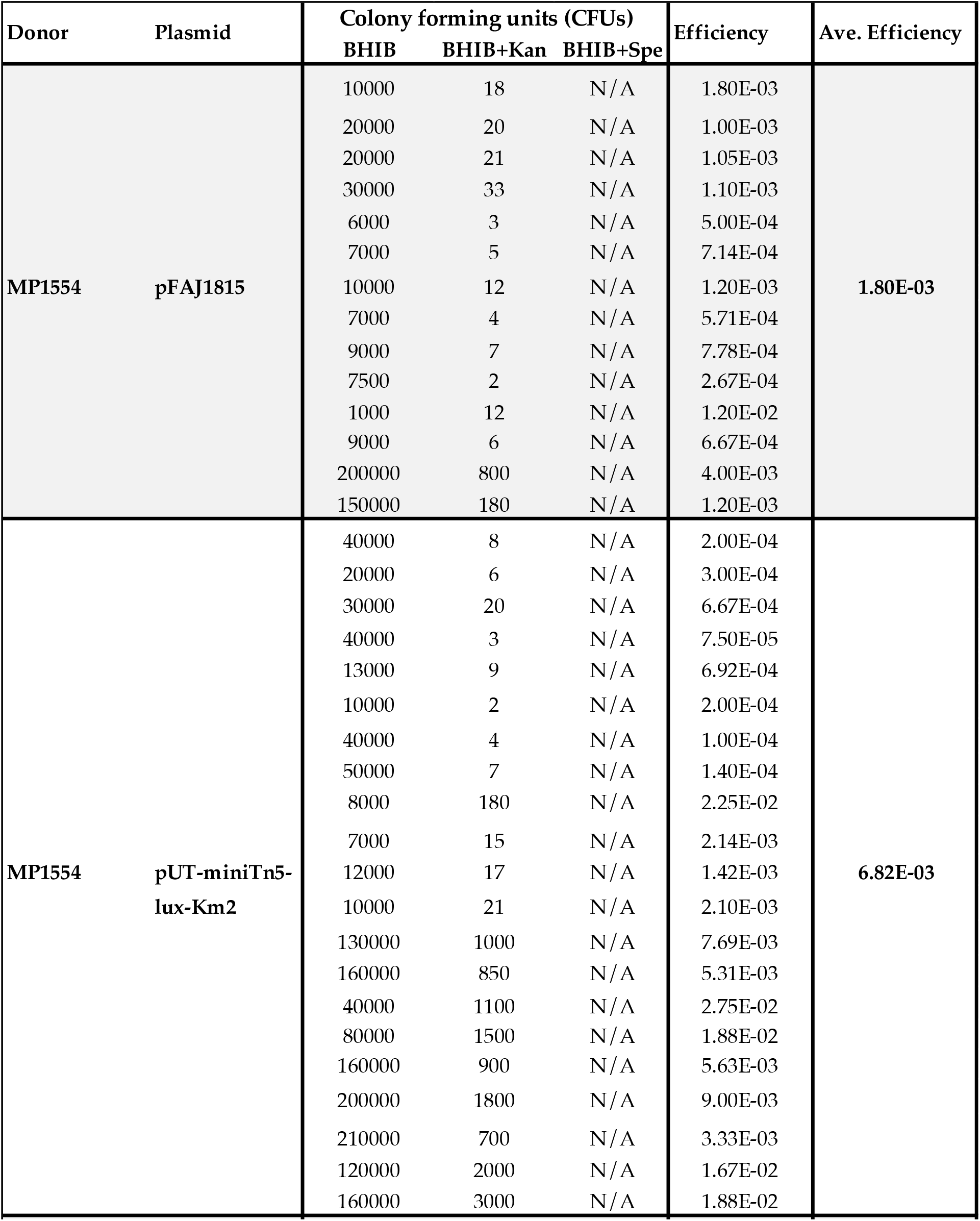
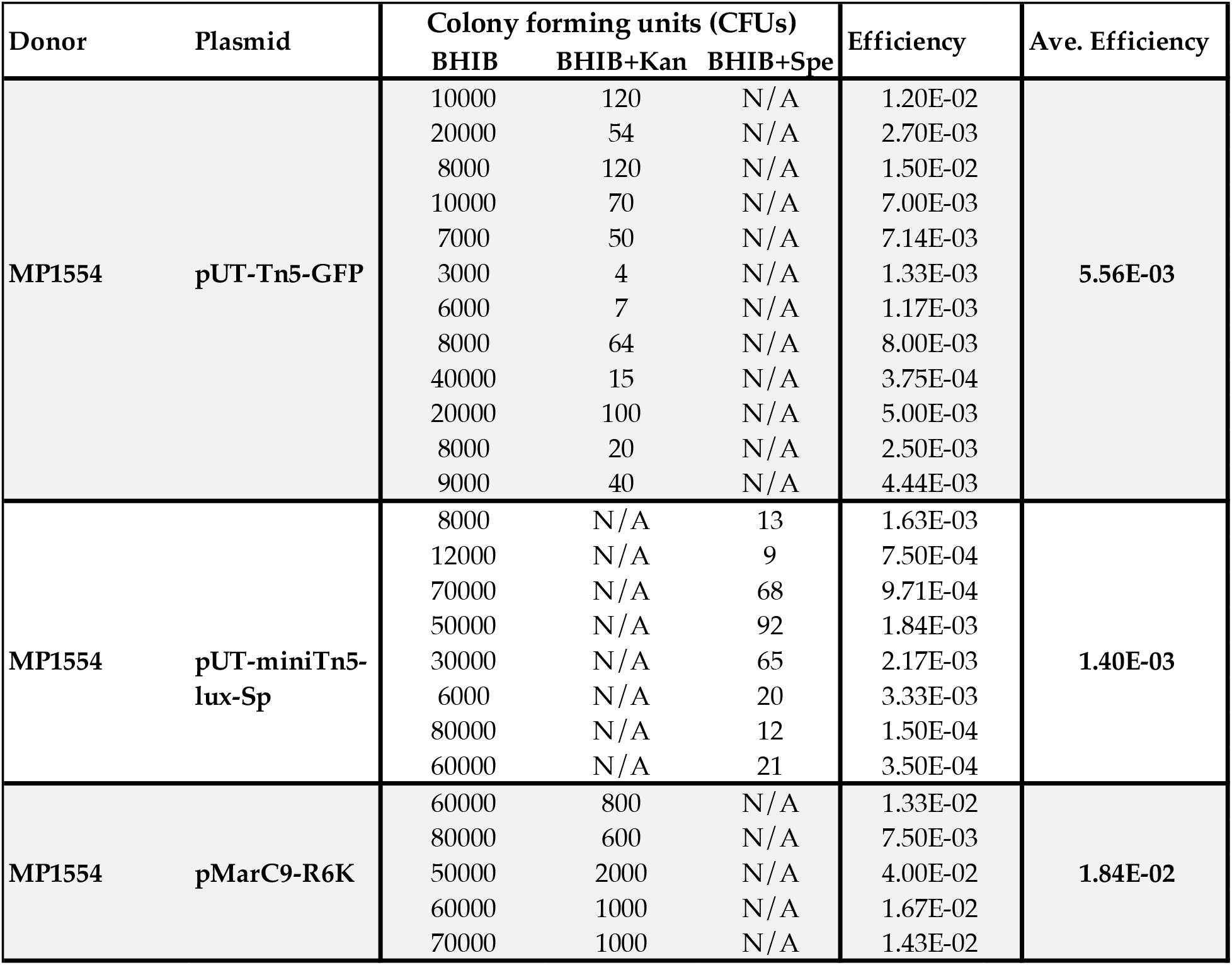
Raw matting experiments

| Donor | Plasmid | Colony forming units (CFUs) |  |  | Efficiency | Ave. Efficiency |
| --- | --- | --- | --- | --- | --- | --- |
|  |  | BHIB | BHIB+Kan | BHIB+Spe |  |  |
| MP1554 | pFAJ1815 | 10000 | 18 | N/A | 1.80E-03 | 1.80E-03 |
|  |  | 20000 | 20 | N/A | 1.00E-03 |  |
|  |  | 20000 | 21 | N/A | 1.05E-03 |  |
|  |  | 30000 | 33 | N/A | 1.10E-03 |  |
|  |  | 6000 | 3 | N/A | 5.00E-04 |  |
|  |  | 7000 | 5 | N/A | 7.14E-04 |  |
|  |  | 10000 | 12 | N/A | 1.20E-03 |  |
|  |  | 7000 | 4 | N/A | 5.71E-04 |  |
|  |  | 9000 | 7 | N/A | 7.78E-04 |  |
|  |  | 7500 | 2 | N/A | 2.67E-04 |  |
|  |  | 1000 | 12 | N/A | 1.20E-02 |  |
|  |  | 9000 | 6 | N/A | 6.67E-04 |  |
|  |  | 200000 | 800 | N/A | 4.00E-03 |  |
|  |  | 150000 | 180 | N/A | 1.20E-03 |  |
| MP1554 | pUT-miniTn5-lux-Km2 | 40000 | 8 | N/A | 2.00E-04 | 6.82E-03 |
|  |  | 20000 | 6 | N/A | 3.00E-04 |  |
|  |  | 30000 | 20 | N/A | 6.67E-04 |  |
|  |  | 40000 | 3 | N/A | 7.50E-05 |  |
|  |  | 13000 | 9 | N/A | 6.92E-04 |  |
|  |  | 10000 | 2 | N/A | 2.00E-04 |  |
|  |  | 40000 | 4 | N/A | 1.00E-04 |  |
|  |  | 50000 | 7 | N/A | 1.40E-04 |  |
|  |  | 8000 | 180 | N/A | 2.25E-02 |  |
|  |  | 7000 | 15 | N/A | 2.14E-03 |  |
|  |  | 12000 | 17 | N/A | 1.42E-03 |  |
|  |  | 10000 | 21 | N/A | 2.10E-03 |  |
|  |  | 130000 | 1000 | N/A | 7.69E-03 |  |
|  |  | 160000 | 850 | N/A | 5.31E-03 |  |
|  |  | 40000 | 1100 | N/A | 2.75E-02 |  |
|  |  | 80000 | 1500 | N/A | 1.88E-02 |  |
|  |  | 160000 | 900 | N/A | 5.63E-03 |  |
|  |  | 200000 | 1800 | N/A | 9.00E-03 |  |
|  |  | 210000 | 700 | N/A | 3.33E-03 |  |
|  |  | 120000 | 2000 | N/A | 1.67E-02 |  |
|  |  | 160000 | 3000 | N/A | 1.88E-02 |  |

**Table S1. Raw matting experiments (cont.)**
| Donor | Plasmid | Colony forming units (CFUs) |  |  | Efficiency | Ave. Efficiency |
| --- | --- | --- | --- | --- | --- | --- |
|  |  | BHIB | BHIB+Kan | BHIB+Spe |  |  |
| MP1554 | pUT-Tn5-GFP | 10000 | 120 | N/A | 1.20E-02 | 5.56E-03 |
|  |  | 20000 | 54 | N/A | 2.70E-03 |  |
|  |  | 8000 | 120 | N/A | 1.50E-02 |  |
|  |  | 10000 | 70 | N/A | 7.00E-03 |  |
|  |  | 7000 | 50 | N/A | 7.14E-03 |  |
|  |  | 3000 | 4 | N/A | 1.33E-03 |  |
|  |  | 6000 | 7 | N/A | 1.17E-03 |  |
|  |  | 8000 | 64 | N/A | 8.00E-03 |  |
|  |  | 40000 | 15 | N/A | 3.75E-04 |  |
|  |  | 20000 | 100 | N/A | 5.00E-03 |  |
|  |  | 8000 | 20 | N/A | 2.50E-03 |  |
|  |  | 9000 | 40 | N/A | 4.44E-03 |  |
| MP1554 | pUT-miniTn5-lux-Sp | 8000 | N/A | 13 | 1.63E-03 | 1.40E-03 |
|  |  | 12000 | N/A | 9 | 7.50E-04 |  |
|  |  | 70000 | N/A | 68 | 9.71E-04 |  |
|  |  | 50000 | N/A | 92 | 1.84E-03 |  |
|  |  | 30000 | N/A | 65 | 2.17E-03 |  |
|  |  | 6000 | N/A | 20 | 3.33E-03 |  |
|  |  | 80000 | N/A | 12 | 1.50E-04 |  |
|  |  | 60000 | N/A | 21 | 3.50E-04 |  |
| MP1554 | pMarC9-R6K | 60000 | 800 | N/A | 1.33E-02 | 1.84E-02 |
|  |  | 80000 | 600 | N/A | 7.50E-03 |  |
|  |  | 50000 | 2000 | N/A | 4.00E-02 |  |
|  |  | 60000 | 1000 | N/A | 1.67E-02 |  |
|  |  | 70000 | 1000 | N/A | 1.43E-02 |  |

**Table S2.**
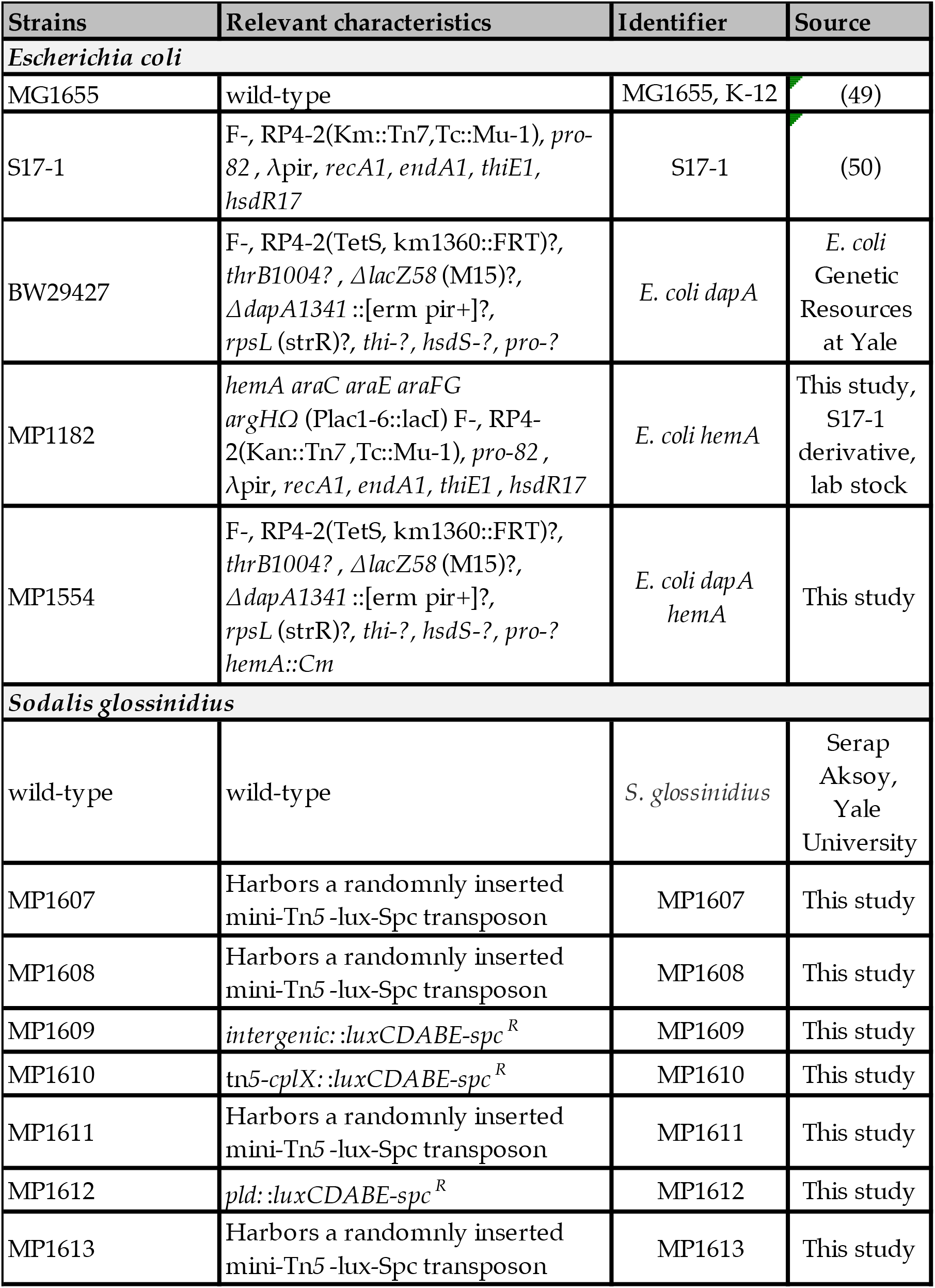
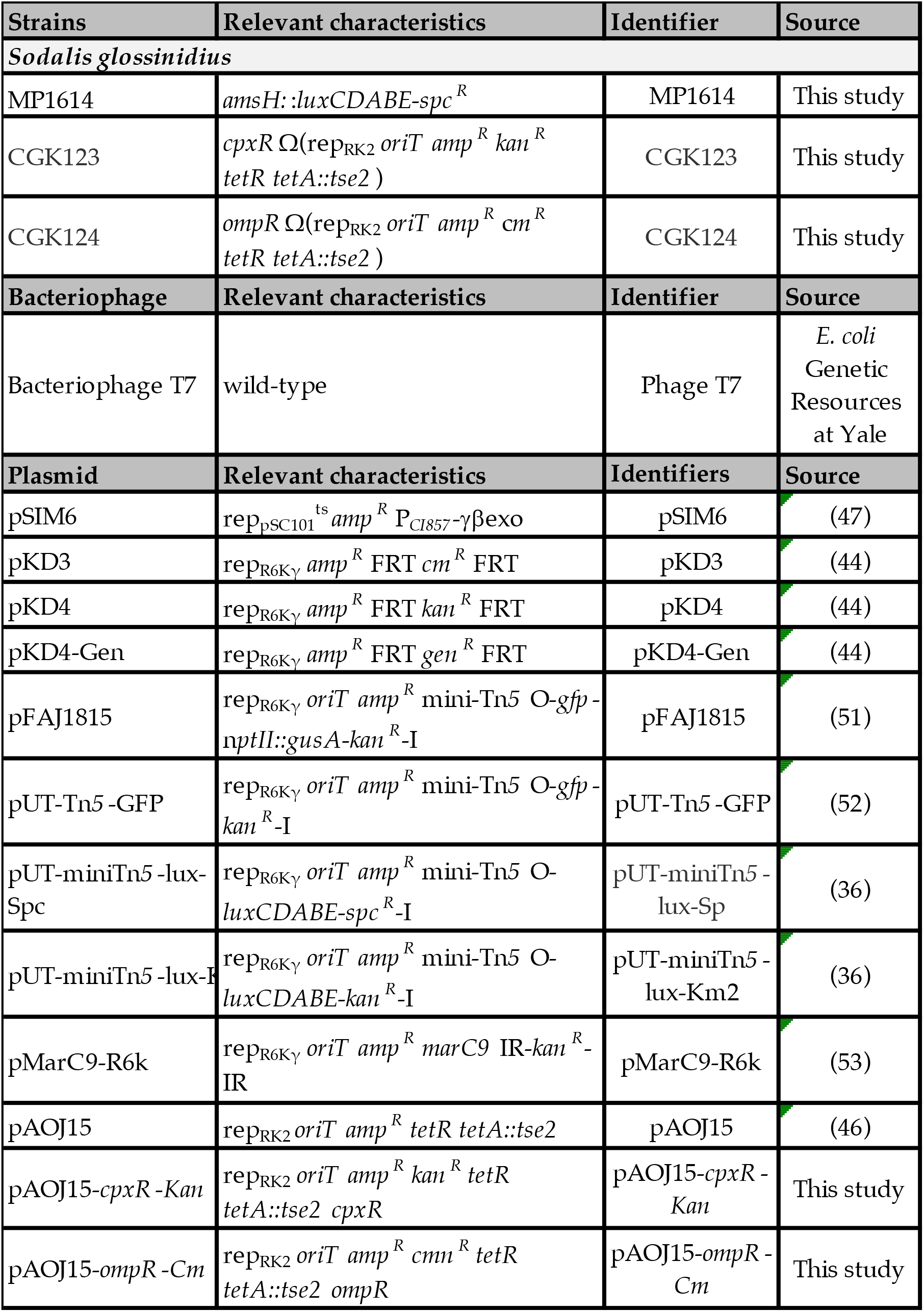
Microbial strains, phages and plasmids used in this study

| Strains | Relevant characteristics | Identifier | Source |
| --- | --- | --- | --- |
| *Escherichia coli* |  |  |  |
| MG1655 | wild-type | MG1655, K-12 | (49) |
| S17-1 | F<sup>-</sup>, RP4-2(Km<sup>+</sup>::Tn7,Tc<sup>+</sup>::Mu-1), *pro*-82, λpir, *recA1*, *endA1*, *thiE1*, *hsdR17* | S17-1 | (50) |
| BW29427 | F<sup>-</sup>, RP4-2(TetS, km1360::FRT)?, *thrB1004*?, Δ*lacZ58* (M15)?, Δ*dapA1341* ::[erm pir+]? , *rpsL* (strR)?, *thi*?, *hsdS*?, *pro*? | *E. coli dapA* | *E. coli* Genetic Resources at Yale |
| MP1182 | *hemA araC araE araFG argH*Ω (Plac1-6::lacI) F<sup>-</sup>, RP4-2(Kan<sup>+</sup>::Tn7,Tc<sup>+</sup>::Mu-1), *pro*-82, λpir, *recA1*, *endA1*, *thiE1*, *hsdR17* | *E. coli hemA* | This study, S17-1 derivative, lab stock |
| MP1554 | F<sup>-</sup>, RP4-2(TetS, km1360::FRT)?, *thrB1004*?, Δ*lacZ58* (M15)?, Δ*dapA1341* ::[erm pir+]? , *rpsL* (strR)?, *thi*?, *hsdS*?, *pro*?, *hemA*::Cm | *E. coli dapA hemA* | This study |
| *Sodalis glossinidius* |  |  |  |
| wild-type | wild-type | *S. glossinidius* | Serap Aksoy, Yale University |
| MP1607 | Harbors a randomly inserted mini-Tn5 -lux-Spc transposon | MP1607 | This study |
| MP1608 | Harbors a randomly inserted mini-Tn5 -lux-Spc transposon | MP1608 | This study |
| MP1609 | *intergenic*::*luxCDABE-spc*<sup>R</sup> | MP1609 | This study |
| MP1610 | *tn5-cplX*::*luxCDABE-spc*<sup>R</sup> | MP1610 | This study |
| MP1611 | Harbors a randomly inserted mini-Tn5 -lux-Spc transposon | MP1611 | This study |
| MP1612 | *pld*::*luxCDABE-spc*<sup>R</sup> | MP1612 | This study |
| MP1613 | Harbors a randomly inserted mini-Tn5 -lux-Spc transposon | MP1613 | This study |

Table S2. Microbial strains, phages and plasmids used in this study (cont.)
| Strains | Relevant characteristics | Identifier | Source |
| --- | --- | --- | --- |
| *Sodalis glossinidius* |  |  |  |
| MP1614 | *amsH::luxCDABE-spc<sup>R</sup>* | MP1614 | This study |
| CGK123 | *cpxR* $\Omega$(*rep<sub>RK2</sub>* *oriT* *amp<sup>R</sup>* *kan<sup>R</sup>* *tetR* *tetA::tse2*) | CGK123 | This study |
| CGK124 | *ompR* $\Omega$(*rep<sub>RK2</sub>* *oriT* *amp<sup>R</sup>* *cm<sup>R</sup>* *tetR* *tetA::tse2*) | CGK124 | This study |
| Bacteriophage | Relevant characteristics | Identifier | Source |
| Bacteriophage T7 | wild-type | Phage T7 | *E. coli*<br>Genetic<br>Resources<br>at Yale |
| Plasmid | Relevant characteristics | Identifiers | Source |
| pSIM6 | *rep<sub>pSC101</sub>* <sup>ts</sup> *amp<sup>R</sup>* *P<sub>CI857</sub>*-γ βexo | pSIM6 | (47) |
| pKD3 | *rep<sub>R6Kγ</sub>* *amp<sup>R</sup>* *FRT* *cm<sup>R</sup>* *FRT* | pKD3 | (44) |
| pKD4 | *rep<sub>R6Kγ</sub>* *amp<sup>R</sup>* *FRT* *kan<sup>R</sup>* *FRT* | pKD4 | (44) |
| pKD4-Gen | *rep<sub>R6Kγ</sub>* *amp<sup>R</sup>* *FRT* *gen<sup>R</sup>* *FRT* | pKD4-Gen | (44) |
| pFAJ1815 | *rep<sub>R6Kγ</sub>* *oriT* *amp<sup>R</sup>* mini-Tn5 O-*gfp* -<br>*nptII::gusA-kan<sup>R</sup>* -I | pFAJ1815 | (51) |
| pUT-Tn5 -GFP | *rep<sub>R6Kγ</sub>* *oriT* *amp<sup>R</sup>* mini-Tn5 O-*gfp* -<br>*kan<sup>R</sup>* -I | pUT-Tn5 -GFP | (52) |
| pUT-miniTn5 -lux-Spc | *rep<sub>R6Kγ</sub>* *oriT* *amp<sup>R</sup>* mini-Tn5 O-<br>*luxCDABE-spc<sup>R</sup>* -I | pUT-miniTn5 -<br>lux-Sp | (36) |
| pUT-miniTn5 -lux-Km2 | *rep<sub>R6Kγ</sub>* *oriT* *amp<sup>R</sup>* mini-Tn5 O-<br>*luxCDABE-kan<sup>R</sup>* -I | pUT-miniTn5 -<br>lux-Km2 | (36) |
| pMarC9-R6k | *rep<sub>R6Kγ</sub>* *oriT* *amp<sup>R</sup>* *marC9* IR-*kan<sup>R</sup>* -<br>IR | pMarC9-R6k | (53) |
| pAOJ15 | *rep<sub>RK2</sub>* *oriT* *amp<sup>R</sup>* *tetR* *tetA::tse2* | pAOJ15 | (46) |
| pAOJ15-*cpxR* -Kan | *rep<sub>RK2</sub>* *oriT* *amp<sup>R</sup>* *kan<sup>R</sup>* *tetR*<br>*tetA::tse2* *cpxR* | pAOJ15-*cpxR* -<br>Kan | This study |
| pAOJ15-*ompR* -Cm | *rep<sub>RK2</sub>* *oriT* *amp<sup>R</sup>* *cmn<sup>R</sup>* *tetR*<br>*tetA::tse2* *ompR* | pAOJ15-*ompR* -<br>Cm | This study |

**Table S3.**
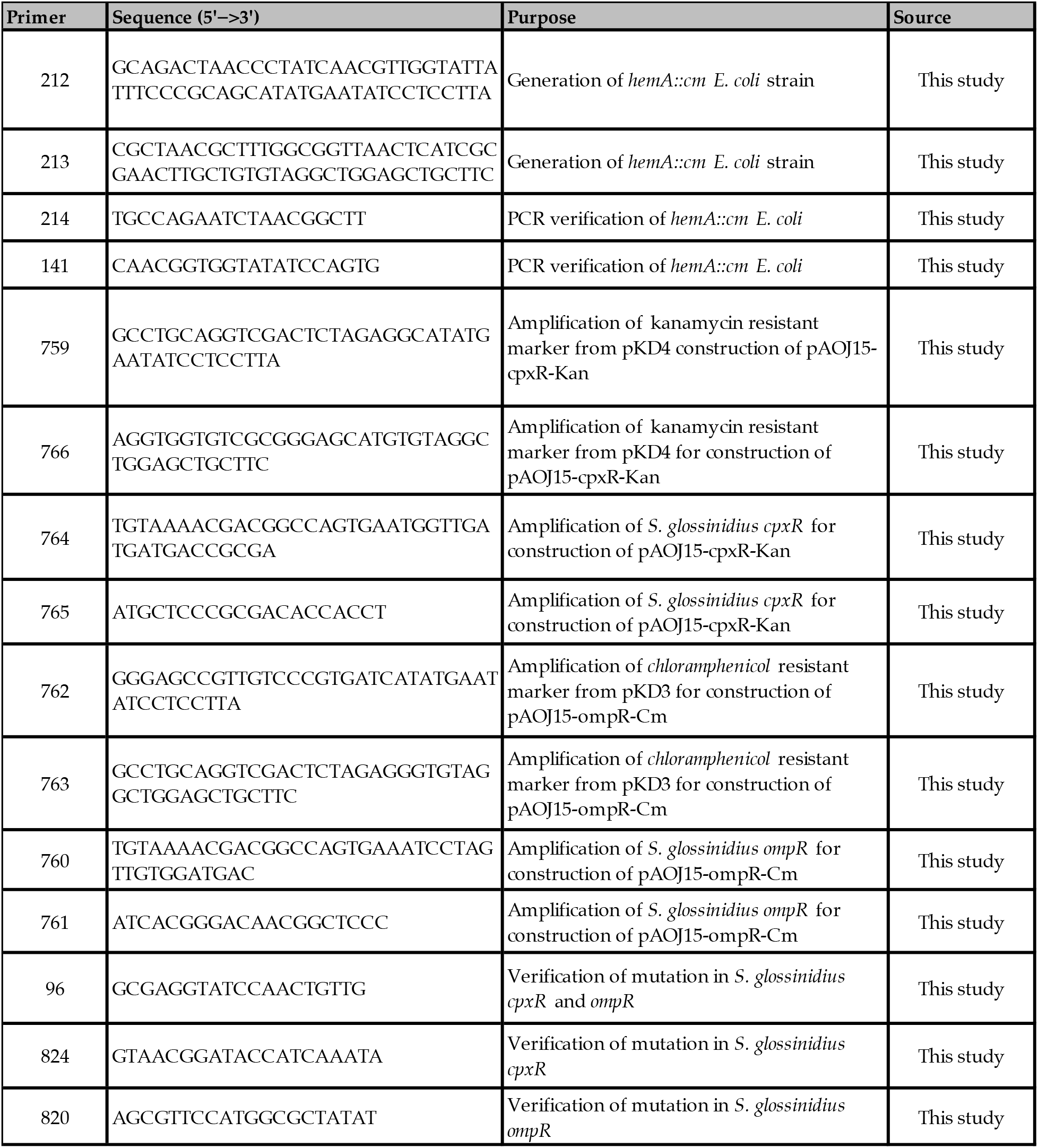
Oligonucleotides sequences used in this study

